# Ecological suicide in microbes

**DOI:** 10.1101/161398

**Authors:** Christoph Ratzke, Jonas Denk, Jeff Gore

**Author notes:** equal contribution (Christoph Ratzke, Jonas Denk). correspondence should be sent to or.

## Abstract

The growth and survival of organisms often depend on interactions between them. In many cases, these interactions are positive and caused by a cooperative modification of the environment, such as the cooperative breakdown of complex nutrients in microbes [1]–[3] or the construction of elaborate architectures in social insects [4]. However, organisms can similarly display negative interactions by changing the environment in ways that are detrimental for them, eg by resource depletion or the production of toxic byproducts [5]. Here we find an extreme type of negative interactions, in which bacteria modify the environmental pH to such a degree that it leads to a rapid extinction of the whole population, a phenomenon we call ecological suicide. Modification of the pH is more pronounced at higher population densities, and thus ecological suicide is more likely with increasing bacterial density. Correspondingly, promoting bacterial growth can drive populations extinct whereas inhibiting bacterial growth by the addition of harmful substances – like antibiotics – can rescue them. Moreover, ecological suicide can cause oscillatory dynamics, even in single-species populations. We find ecological suicide in a wide variety of microbes, suggesting that it could play a significant role in microbial ecology and evolution.

## Once sentence summary

Bacterial populations can cause their own extinction by collectively deteriorating the environment

## Main

Microbes depend on their environment but also modify it. An especially important environmental parameter for microbial growth is the pH, since protein and lipid membrane stability depend strongly on it [6], [7]. Microbes have a species-dependent pH optimum at which they grow best [8], [9], and environmental pH away from this optimum either inhibits growth or can even cause cell death [10], [11]. At the same time, bacteria change the environmental pH by their metabolic activities [10], [12]. In this way, microbes can potentially induce pH values that are detrimental for their own growth and thus harm themselves.

The soil bacterium *Paenibacillus sp*. (most similar to *Paenibacillus tundrae*, for more information about this strain see Supplementary Information) can grow in a medium that contains 10g/L glucose as the main carbon source, in addition to a small amount of complex nutrients (see Methods for details). Starting from neutral pH we measured a strong acidification of the environment to a pH of around 4 during bacterial growth (Fig. 1a). Upon reaching this low pH, the bacteria suddenly start to die, resulting in a non-monotonic growth curve (Fig. 1a). Indeed, after 24 hours of incubation, we find that there are no viable cells in the culture (as measured by colony forming units). We call this rapid population extinction due to environmental modification “ecological suicide”.

**Fig. 1:**
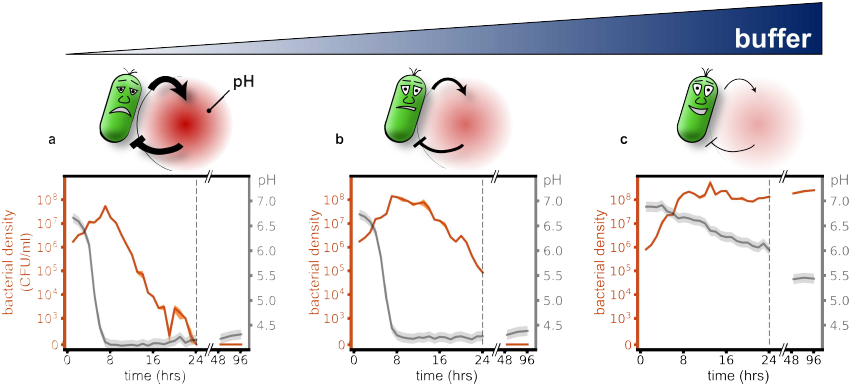
Microbial acidification can cause ecological suicide. Paenibacillus sp. was grown in well- mixed batch culture in media containing 10g/L glucose as main carbon source and minor amounts of complex nutrients (see methods). **(a)** At low buffer concentrations (10mM phosphate) initially growing bacteria change the pH of the medium so drastically that they cause their own extinction. **(b-c)** Adding increasing amounts of buffer (14 and 100mM phosphate) tempers the acidification, and finally allows for survival of the bacteria.

The correlation between the drop of pH and the onset of death suggests that the bacteria themselves may be responsible for their eventual extinction by lowering the pH into regions in which they cannot survive. To test this idea, we added buffer to the medium to temper the pH change. The buffer indeed slows down the death process (Fig. 1b) and prevents it completely at sufficiently high concentrations (Fig. 1c). Thus, it is the pH change that causes the death of the bacteria. These results show that initially flourishing bacterial populations can corrupt their environment and thus cause their own extinction. The pH change resembles a 'public bad' that is collectively produced and harms all members of the populations. This phenomenology can be recapitulated by a simple mathematical description based on negative feedback of the bacteria and the environmental pH (Supplementary Discussion and Fig. S6).

Since bacteria collectively change the pH, higher bacterial densities can deteriorate the environment more strongly and thus expedite ecological suicide. We tested this idea experimentally by measuring the fold growth within 24 hours for different initial bacterial densities and different buffer concentrations. At low buffer concentrations, the bacteria die by ecological suicide independent of their initial density, whereas at high buffer concentrations they always survive (Fig. 2a). At intermediate buffer concentrations, however, survival becomes density-dependent (Fig. 2a). For high initial cell densities, the bacteria die within 24 hours, but below a critical initial density, the bacteria grow and survive. The fitness of the bacteria thus decreases dramatically with increasing cell density. This aspect of ecological suicide is thus opposite of the well-known Allee effect, where fitness increases with population density [13]–[15].

**Fig. 2:**
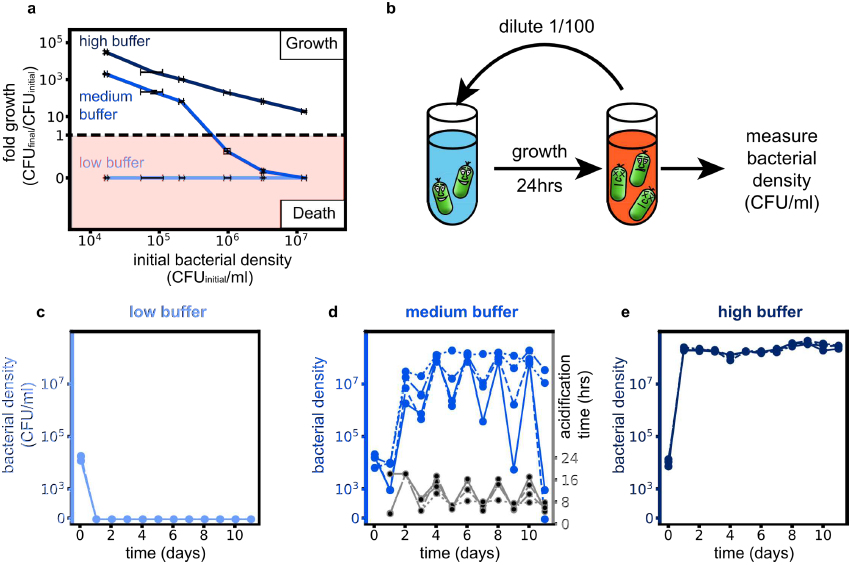
Ecological suicide can cause oscillations in the population size over time. **(a)** At low buffer concentration (10mM phosphate) the bacteria commit ecologic suicide and at high buffer concentration (100mM phosphate) the bacteria grow, in both cases independent of their initial density. However, in moderate buffer concentration (26mM phosphate) the bacteria die at high starting densities and grow at low starting densities. The fold growth at high buffer concentration decreases for increasing initial bacterial densities since the final bacterial density equals the carrying capacity and is thus constant. **(b)** To explore long time growth dynamics the bacteria were grown in a daily dilution scheme with 24h of incubation in well mixed conditions followed by a 1/100× dilution into fresh media. **(c and e)**. At low (10mM phosphate) and high (100mM phosphate) buffer conditions the bacteria either die on the first day or grow to saturation every day. **(d)** However, at medium buffer conditions we measure oscillatory dynamics of the bacterial density. This is accompanied by oscillations in the time the bacteria need to acidify the environment (acidification time, Fig. S5). The exact type of oscillatory dynamics depends on the slope and shape of the curve in **(a)**, as discussed in more detail in the supplement. The 4 blue lines in **(c-e)** (solid, dashed, dotted, dashed-dotted) show different replicates.

What does this growth behavior mean for the long-term growth dynamics of a population such as occurs in growth with daily dilution into fresh media? Figure 2a shows how the bacterial density after one day of growth depends on the initial bacterial density. For intermediate buffering the bacteria die for high initial densities but grow for low initial densities. This may cause oscillatory dynamics, since high bacterial densities cause low densities on the next day and vice versa. Indeed, this intuitive prediction is fully supported by a mathematical description based on negative feedback of the bacteria and the environmental pH alone, which shows a bifurcation of the end-of-the-day bacteria densities upon changing the buffer concentration (see Supplementary Information and Fig. S6 and S7). To test this prediction, we cultivated the bacteria in batch culture with a daily dilution of the culture 1/100× into fresh media. As expected from Fig. 2a, for low buffering the bacteria go extinct on the first day and for high buffering they grow up to the same saturated density each day (Fig. 2c, e, Fig. S3 and S5a). For intermediate buffering, however, the bacteria show oscillatory dynamics as predicted by our model (Fig. S7 and S8). The oscillations of the populations are accompanied by oscillations of the time at which the pH drops each day (acidification time, Fig. 2d and Fig. S2 and S5b), which again shows the connection between pH change and ecological suicide. Ecological suicide caused by environmental deterioration therefore can drive oscillatory dynamics even in populations consisting of just one species.

We have seen that low bacterial densities lead to less deterioration of the environment and thus a less deadly effect on the bacteria. Therefore, effects that hinder bacterial growth by harming the bacteria may be able to save the population from ecological suicide. A first hint in this direction is given by changing the glucose concentration. While one would naively expect that higher glucose concentrations are beneficial, in the presence of ecological suicide, the opposite is the case (Fig 3a). At low glucose concentrations, the bacteria grow to lower densities, which hardly changes the pH and therefore allows the bacteria to survive. At high glucose concentrations, bacterial growth causes environmental acidification and thus ecological suicide. The bacterial population is therefore only able to survive in nutrient-poor conditions. Moreover Fig. 3a shows that ecological suicide can be observed even at rather low nutrient concentrations of around 0.2% glucose.

**Fig. 3:**
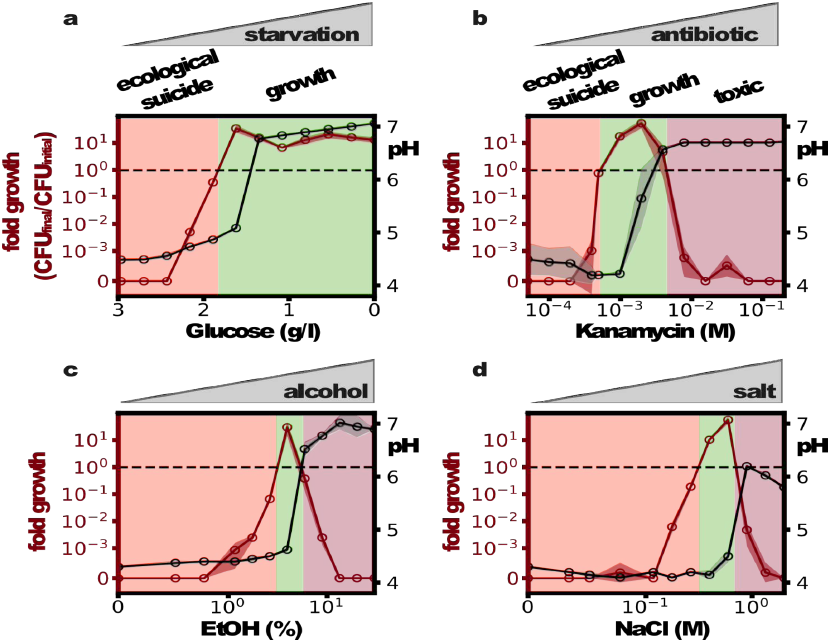
Harming the bacteria saves the population. **(a)** Reducing sugar concentration prevents ecological suicide. At moderate concentrations, the addition of bactericidal substances like antibiotics **(b)**, alcohol **(c)** or high amounts of sodium chloride **(d)** can save the population from ecological suicide.

To explore the idea that poor environments in general can save the bacterial population, we measured the growth and survival of bacteria grown in the presence of the antibiotic kanamycin, ethanol, and salt. Although these substances are quite different, they all inhibit bacterial growth and lead to similar profiles of population survival as a function of the concentration of the inhibiting substance (Fig. 3 b-d). In the absence of the harmful substances, the bacteria lower the pH to the point of extinction. At high concentrations, the harmful substances kill the bacteria. However, at intermediate concentrations, the bacteria can grow and survive. This leads to the paradoxical situation that substances that are normally used to kill bacteria in medicine (antibiotics) or food preservation (salt, ethanol) are able to save bacteria and allow their growth.

The effect of ecological suicide is surprising and has paradoxical consequences. However, the question arises: How common is ecological suicide in bacteria? To investigate this question, we incubated 119 bacterial soil isolates from a wide taxonomic range in the presence of glucose as a carbon source and urea as a nitrogen source. Glucose can be converted to organic acids and acidify the medium, whereas urea can be converted by many bacteria into ammonia and alkalize the environment. From these 119 strains, the 22 strongest pH modifiers (either in acidic or alkaline directions) were tested for the presence of ecological suicide by measuring the fold growth in 24h at low and high buffer concentrations (Fig. 4a). Indeed, around 25% of the strains suffered ecological suicide and were unable to survive at low buffer concentrations yet could be saved by more buffering (Fig. 4b). Another 20% grew better at high than low buffer, suggesting a self-inhibiting but non-deadly effect of the pH. Finally, one species even changed the pH in ways that supported its own growth (an effect discussed in more detail in a separate manuscript [10]). These results show that ecological suicide is not an exotic effect but appears rather often and may therefore play a major role in limiting the environments that microbial species can survive in.

**Fig. 4:**
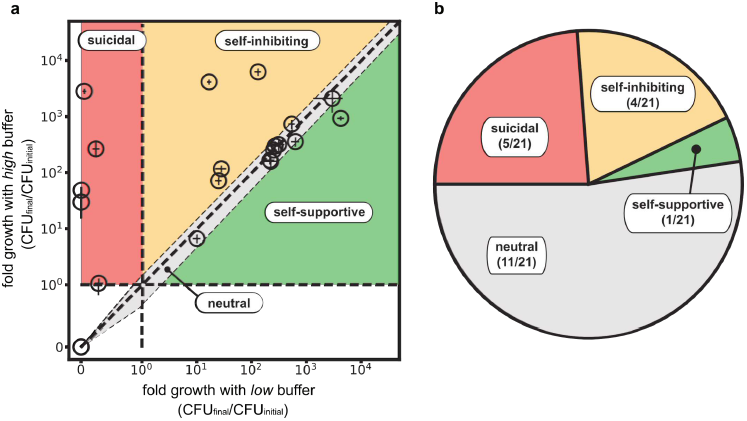
Ecological suicide is a common phenomenon in microbes. 21 bacteria that strongly modified the pH were tested for ecological suicide by growing them on a medium containing 10g/L glucose and 8g/L urea at low buffer (10mM Phosphate) and high buffer (100mM Phosphate) conditions. Bacteria that die at low buffer but grow at high buffer concentrations were counted as ecological suicide (suicidal, 5). Bacteria that grow slower at low buffer than high buffer conditions are called self-inhibiting (4). Bacteria that grow in similar ways at low and high buffer (growth in one buffer condition is between equal and 1.5 fold relative to growth in the other condition) were called neutral (11) and bacteria that grow better with low than with high buffer are called self-supportive (1).

We demonstrated that microbes are able to cause their own extinction by deteriorating the environment, a process we call ecological suicide. Several cases are described where microbial populations experience a slow decline after reaching saturation [16], [17]. However, this decline is usually very slow compared to the growth rate and does not cause sudden population extinction. In ecological suicide, however, the population does not even reach saturation; instead, the bacteria switch immediately from a growth into a death phase (Fig 1a). A notable exception are quorum sensing deficient mutants of several *Burkholderia specie*s that show a type of ecological suicide [18], whereas in the wildtype strains quorum sensing mediates a change of metabolism that avoids ecological suicide. This shows that bacteria can possess mechanisms that actively counteract ecological suicide [18]–[20].

A phenomenon similar but not identical to ecological suicide is population overshoot, which is often connected to overexploitation of natural resources and has been proposed in several macro- organisms [21]–[23], but it is mostly discussed in humans that over-exploit the environment [24]–[26]. Several ancient civilizations are suspected to have collapsed by overexploitation of natural resources [27]–[29]. Upon overshoot, a population exceeds the long-term carrying capacity of its ecosystem, followed by a drop of the population below the carrying capacity which usually does not lead to extinction of the population but is followed by recovery at a lower density [21], [26]. However, in our case of ecological suicide, the carrying capacity of the ecosystem is changed to zero – the bacteria produce a deadly environment and go extinct without recovery, which marks ecological suicide as an extreme version of population overshoot.

In daily dilutions, ecological suicide can result in oscillatory behavior. Oscillations in ecology have been intensely studied, often as a consequence of species interactions [30], [31]; in our system the second species is replaced by the pH value, resulting in a situation where interactions between one species and its environment drive the oscillations. In a similar way modifying and reacting to the environment have recently been described to cause metabolic oscillations in yeast [32] or expanding waves in microbial biofilms [33].

In view of the high frequency of ecological suicide that we observed in natural isolates of soil bacteria, this effect may have a broad impact on microbial ecosystems in terms of microbial interactions and biodiversity [10]. In our case, the ecological suicide was mediated by the pH, but changing any environmental parameter, like oxygen levels or metabolite concentrations in self-harming ways may cause similar outcomes.

Our findings raise the question of how such self-inflicted death of microbes can exist in nature. We speculate that although ecological suicide is detrimental for the population it may be evolutionary beneficial for the individual bacterium. A fast metabolism of glucose may harm and even kill the population but benefits the individual compared to an individual that takes the burden of slower glucose metabolism to save the population. The phenomena of ecological suicide could therefore be an end-product of evolutionary suicide [34]. Future work will explore the evolutionary origin of ecological suicide as well as the consequences of this phenomenon for the ecology and evolution of microbes.

